# Zebrafish GPR161 Contributes to Basal Hedgehog Repression in a Tissue-specific Manner

**DOI:** 10.1101/616482

**Authors:** Philipp Tschaikner, Dominik Regele, Willi Salvenmoser, Stephan Geley, Eduard Stefan, Pia Aanstad

## Abstract

Hedgehog (Hh) ligands act as morphogens to direct patterning and proliferation during embryonic development. Protein kinase A (PKA) is a central negative regulator of Hh signalling, and in the absence of Hh ligands, PKA activity prevents inappropriate expression of Hh target genes. The G_αs_- coupled receptor Gpr161 contributes to the basal Hh repression machinery by activating PKA, although the extent of this contribution is unclear. Here we show that loss of Gpr161 in zebrafish leads to constitutive activation of low-, but not high-level Hh target gene expression in the neural tube. In contrast, in the myotome, both high- and low-level Hh signalling is constitutively activated in the absence of Gpr161 function. Our results suggest that the relative contribution of Gpr161 to basal repression of Hh signalling is tissue-specific. Distinct combinations of G-protein-coupled receptors may allow the fine-tuning of PKA activity to ensure the appropriate sensitivity to Hh across different tissues.

## Introduction

The Hh signalling pathway is a key regulator of cell fate specification and proliferation during embryonic development, and plays important roles in adult tissue homeostasis (Briscoe and Thérond, 2013; Ingham et al., 2011). Dysregulation of Hh signalling can lead to the formation of common and severe forms of human cancers such as basal cell carcinoma and medulloblastoma (Jiang and Hui, 2008; Raleigh and Reiter, 2019).

When Hh ligands bind their receptor Patched (Ptch), the inhibition of Smo by Ptch is alleviated, and Smo localises to the primary cilium (Corbit et al., 2005), where it activates downstream signalling to regulate the activity of the bifunctional Gli transcription factors.

Hh ligands act as morphogens, and the transcriptional outcome of Hh signalling is determined by the balance between repressor and activator forms of the Gli transcription factors. This balance is controlled by the activity of PKA and other kinases (Hui and Angers, 2011; Niewiadomski et al., 2019). In the absence of Hh, the basal Hh repression machinery is thought to maintain a high level of PKA activity. PKA phosphorylates the Gli proteins and primes them for further phosphorylation and proteolytic cleavage, to yield truncated forms that act as transcriptional repressors (GliR) (Niewiadomski et al., 2014; Pan et al., 2009; Wang et al., 2000). In addition, PKA also plays a role in restricting the activity of full length Gli (GliA) by promoting its association with Sufu (Humke et al., 2010; Marks and Kalderon, 2011). Low levels of exposure to Hh ligands blocks the formation of GliR, whereas high levels of Hh exposure is required for the formation of the activator forms of Gli. This is thought to be controlled through a cluster of six PKA target residues in Gli, with distinct phosphorylation patterns regulating the formation of repressor and activator forms (Niewiadomski et al., 2014). This rheostat mechanism ensures that the level of Gli transcriptional activity corresponds to the level of PKA activity, which in turn must be controlled by the level of Smo activity and Hh ligand exposure (Niewiadomski et al., 2019, 2014). Consistent with this, a complete loss of PKA activity leads to constitutive (Smo-independent) maximal Hh signalling, whereas constitutive activation of PKA abolishes all Hh-dependent transcription (Hammerschmidt et al., 1996; Tuson et al., 2011; Zhao et al., 2016).

A central and long-standing question in Hh signalling regards the nature and regulation of the basal repression machinery and the mechanism of regulation. In *Drosophila*, Smo has been shown to regulate PKA activity directly by activating G_αi_-proteins to modulate cAMP levels (Ogden et al., 2008). Although vertebrate Smo can also couple to G_αi_ (Riobo et al., 2006), it is clear that G_αi_ is not required for all aspects of vertebrate Hh signalling (Ayers and Thérond, 2010), raising the question which other mechanisms contribute to the regulation of PKA.

The murine orphan G-protein coupled receptor (GPCR) Gpr161 contributes to basal Hh repression by activating G_αs_ and consequently PKA (Mukhopadhyay et al., 2013). In the absence of Hh ligands, Gpr161 localises to the primary cilia, and is removed from the cilia upon activation of Smo (Mukhopadhyay et al., 2013; Pal et al., 2016). These results suggest that Gpr161 maintains PKA activity in the cilium in the absence of Hh, and that the ciliary exit of Gpr161 is required for Hh signalling and the reduction of PKA activity (Mukhopadhyay et al., 2013; Pal et al., 2016). However, Gpr161 has also been shown to be a substrate of PKA, and can act as an A kinase anchoring protein (Bachmann et al., 2016; Torres-Quesada et al., 2017). Thus, the exact molecular mechanisms that regulate Gpr161 activity in the context of Hh signalling remain unclear.

Gpr161 mutants display severe developmental malformations, including craniofacial defects and ventralisation of the neural tube, that were independent of Smo function, suggesting that Gpr161 causes constitutive activation of downstream Hh signal transduction (Mukhopadhyay et al., 2013). However, the neural tube of Gpr161 mutants was less severely ventralised than that observed in embryos completely lacking PKA activity (Tuson et al., 2011). In neural progenitor cells (NPCs), Gpr161 was found to be epistatic to Smo only for low level signalling, while expression of high level targets such as Nkx2.2 and FoxA2 still depended on Smo function (Pusapati et al., 2018), suggesting that in the neural tube, Gpr161 plays an important role in controlling basal and low level Hh signalling activity. In murine NIH 3T3 fibroblasts, loss of Gpr161 does not affect basal repression, although the mutant cells displayed an increased sensitivity to Hh ligands. Taken together, these results suggest that additional G_αs_-coupled receptors may be involved in maintaining PKA activity in the absence of Hh ligands (Pusapati et al., 2018). Supporting this, several studies have identified additional GPCRs that regulate Hh signalling in parallel or downstream of Smo (Klein et al., 2001; Singh et al., 2015; Stückemann et al., 2012).

To facilitate the study of Gpr161 in Hh signalling during development, we have generated zebrafish *gpr161* mutants, and show that Gpr161 is an important negative modulator of Hh signalling in zebrafish embryos. We find that in the zebrafish neural tube, Gpr161 is epistatic to Smo for low-level Hh targets, however the activation of high-level targets depends on Smo activity in *gpr161* mutants. Interestingly, in the myotome, both high and low levels of Hh signalling are independent of Smo function, suggesting that several GPCRs may be involved in regulating PKA activity during Hh signalling, and that the level of contribution of Gpr161 to basal repression of Hh signalling is tissue-specific.

## Results

### Gpr161 is an evolutionary conserved GPCR with two paralogs in zebrafish

The zebrafish genome contains two conserved paralogs of Gpr161, Gpr161a and Gpr161b, with 71% sequence identity and 84% sequence similarity between each other, and more than 70% similarity to the murine Gpr161 protein (Figure 1A, Figure 1 Supplement 1). Expression analysis using qRT-PCR showed that both transcripts are expressed during embryonic development, but while *gpr161b* is maternally provided, *gpr161a* expression is first detected at 9 hours post fertilisation (hpf) (Figure 1B). In mouse, Gpr161 localises to primary cilia, and this localisation has been proposed to be important for its role in modulating the Hh signalling pathway (Mukhopadhyay et al., 2013; Pal et al., 2016; Shimada et al., 2018). To test whether zebrafish Gpr161 also localise to primary cilia, we injected mRNA of Myc-tagged versions of Gpr161a and Gpr161b into one-cell stage zebrafish embryos. Both Gpr161a and Gpr161b were readily detected at primary cilia in gastrula stage zebrafish embryos (Figure 1C).

**Figure 1.**
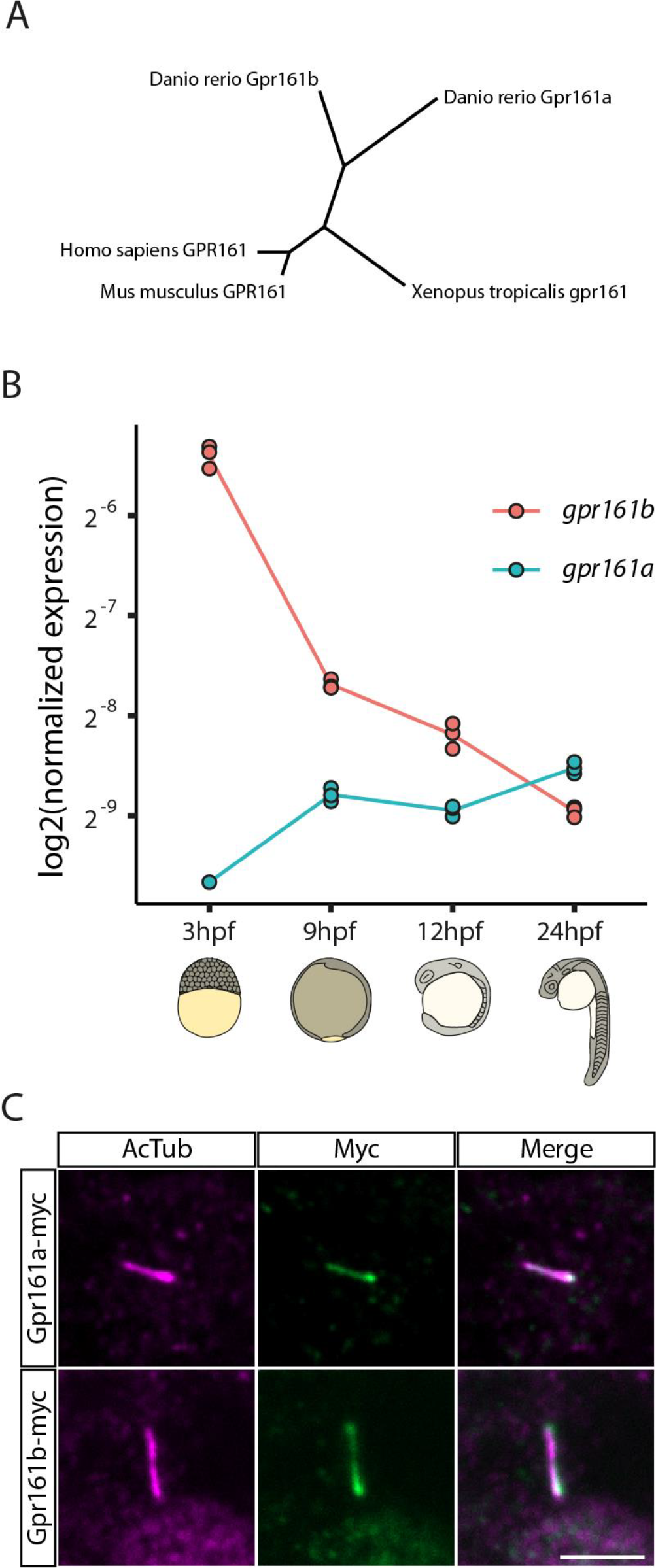
Gpr161 is a conserved ciliary GPCR. **(A)** Un-rooted cladogram showing the relation between Gpr161 protein sequences of selected organisms. **(B)** *gpr161a* and *gpr161b* transcript levels at different stages of development analysed in whole embryo lysates in triplicates using RT-qPCR. **(C)** Wildtype embryos were injected with *gpr161a-myc* or *gpr161b-myc* mRNA at one-cell stage and fixed at 9 hpf before immunostaining for acetylated tubulin (AcTub; purple), a marker for the ciliary axoneme and Myc (green) (scale bar: 5µm).

### Gpr161a and Gpr161b are functionally redundant but essential during zebrafish embryonic development

To investigate the functional roles of Gpr161a and Gpr161b during zebrafish development, we used CRISPR/cas9 to generate mutant alleles for each gene, *gpr161a*^*ml200*^, which carries a 6bp insertion in the second coding exon of *gpr161a*, and *gpr161b*^*ml201*^, harboring an 8bp deletion in the second coding exon of *gpr161b*. Both alleles introduce premature stop codons within the 7-transmembrane-domain region of the respective proteins, and are predicted to be functionally inactive (Figure 1 Supplement 2). Animals homozygous for either *gpr161a*^*ml200*^ or *gpr161b*^*ml201*^ alone, or animals lacking three of the four *gpr161* alleles, showed no effect either on embryonic development or in adult viability and fertility. In contrast, *gpr161a; gpr161b* double zygotic homozygous mutant embryos showed clear morphological phenotypes by 24 hpf, suggesting that Gpr161a and Gpr161b act in a functionally redundant manner (Figure 2A). To determine the contribution of maternal Gpr161b, we generated embryos from incrosses of *gpr161b*^−/−^; *gpr161a*^+/−^ animals. Quantitative analysis of *gpr161a* expression at the 2 cell stage of these *MZgpr161b; gpr161a* mutant embryos showed that a complete loss of Gpr161b did not result in a compensatory maternal upregulation of Gpr161a (Figure 2 Supplement 1). We conclude that *MZgpr161b*^−/−^; *gpr161a*^−/−^ mutant embryos are likely to represent a complete loss of function of zebrafish Gpr161, and refer to these mutants as *gpr161* mutants below.

**Figure 2.**
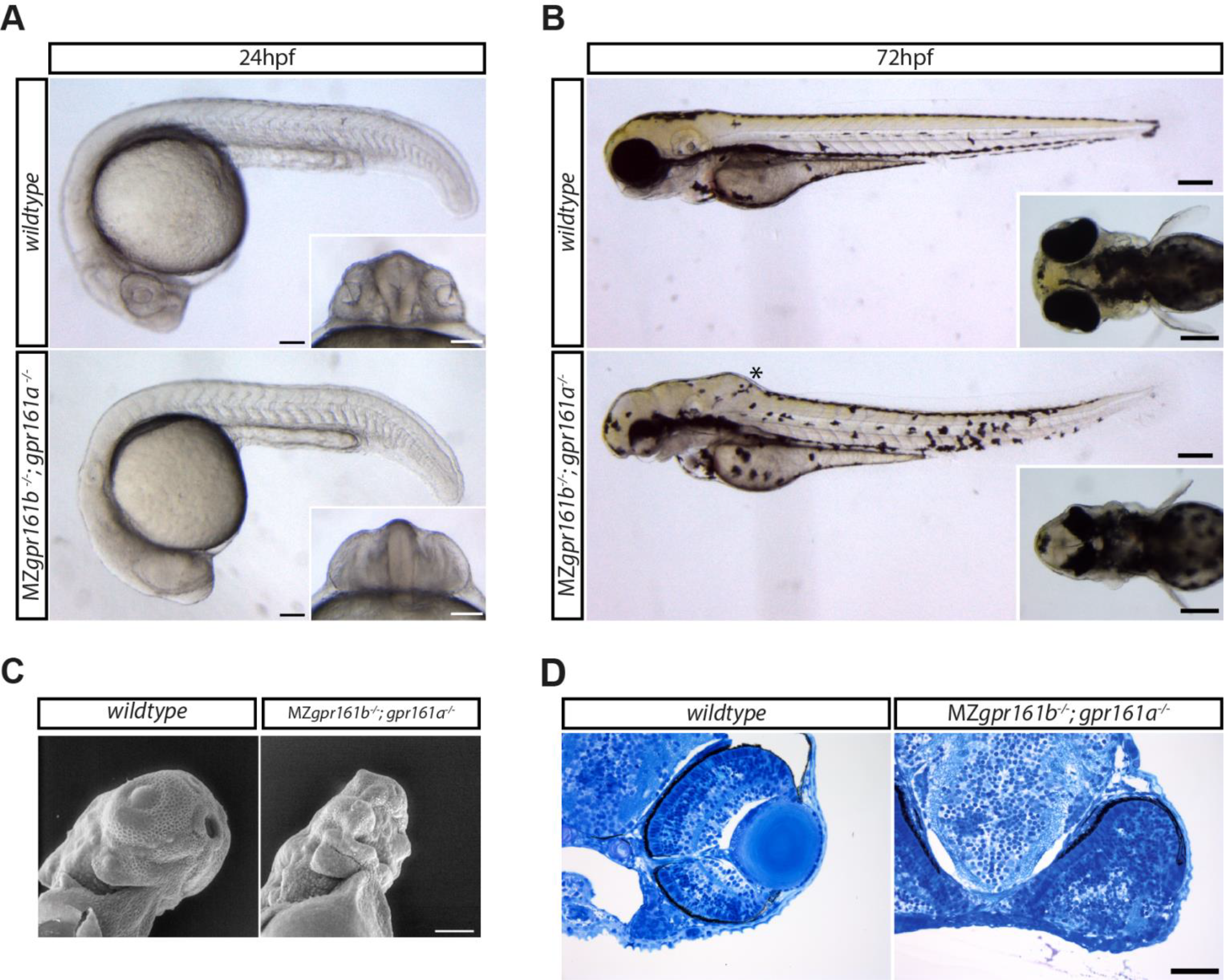
Gpr161 is essential during embryonic development. **(A)** Lateral view of wildtype and MZ*gpr161b*^−/−^; *gpr161a*^−/−^ embryos at 24 hpf (scale bars: 100µm); Insets: ventral view of the developing eyes. **(B)** Wildtype and MZ*gpr161b*^−/−^; *gpr161a*^−/−^ embryos at 72 hpf; Asterisk indicates swollen hindbrain (scale bars: 200µm); Insets: dorsal view of the head **(C)** Ventrolateral-view of the craniofacial region of wildtype and MZ*gpr161b*^−/−^; *gpr161a*^−/−^ embryos at 72 hpf taken by scanning electron microscopy (scale bar: 100µm). **(D)** Transverse semi-thin sections of the eye in wildtype and MZ*gpr161b*^−/−^; *gpr161a*^−/−^ embryos fixed at 72 hpf (Richardson staining, scale bars: 50µm).

At 24hpf, *gpr161* mutant embryos display several developmental abnormalities, including malformed eyes lacking any obvious lens or retinal structure (Figure 2A). At this stage of development, wildtype embryos display chevron shaped somites, while the somites of *gpr161* mutants have a more obtuse angle (Figure 2A). These phenotypes, which are only present in the double mutant line, but not in MZ*gpr161b*^−/−^; *gpr161a*^+/−^ or *gpr161b*^+/−^; *gpr161a*^−/−^ embryos, are similar to those observed in *ptch1*^−/−^; *ptch2*^−/−^ double mutant embryos (Koudijs et al., 2008). In contrast to *ptch1*^−/−^; *ptch2*^−/−^ double mutant embryos which lack eyes (Koudijs et al., 2008), a rudimentary eye can be identified in *gpr161* mutants at 72 hpf (Figure 2B).

At this stage, the retina of wild type embryos is organised into six evolutionarily conserved layers: the pigmented epithelium, the photoreceptor cell layer, the outer plexiform layer, the inner nuclear layer, the inner plexiform layer, and the ganglion cell layer (Schmitt and Dowling, 1999). Semi-thin sectioning revealed that while eye morphogenesis was abnormal, remnants of all six layers were clearly identified by morphology (Figure 2D, Figure 2 Supplement 2A). However, these layers are not well separated and the optic cup is partially folded (Figure 2 Supplement 2B). Additionally, remnants of a forming lens can be found in *gpr161* mutant embryos (Figure 2 Supplement 2A). A striking phenotype of the *gpr161* mutants is the complete loss of jaw structures (Figure 2B), as demonstrated by scanning electron microscopy (SEM) imaging, which revealed that *gpr161* mutant embryos display an open pharynx with a complete lack of all jaw structures (Figure 2C). While *ptch1*^−/−^; *ptch2*^−/−^ double mutant embryos were reported to lack all olfactory structures (Koudijs et al., 2008), SEM imaging revealed that in the *gpr161* mutants the olfactory pits are present, though severely reduced in size (Figure 2C).

Zebrafish embryos with a strong Hh gain-of-function phenotype, such as the *ptch1*^−/−^; *ptch2*^−/−^ double mutants, also display defects in the development of the otic vesicles (Koudijs et al., 2008). The *gpr161* mutant embryos exhibited smaller otic vesicles compared to wild type embryos (Figure 2B). Serial sections of the otic vesicles revealed the absence of the dorsolateral septum (Figure 2 Supplement 2C). The combination of developmental defects in ocular and otic structures is commonly seen in mutants of negative regulators of Hh signalling, such as *sufu*, *ptch1* and *hhip* (Whitfield et al., 1996), and suggests that Gpr161 acts to negatively regulate Hh signalling also in zebrafish.

Gpr161 mutant mice do not form limb buds (Hwang et al., 2018; Mukhopadhyay et al., 2013) and *ptch1*^−/−^; *ptch2*^−/−^ double mutant zebrafish embryos lack pectoral fin buds (Koudijs et al., 2008). In contrast to the requirement for Gpr161 in murine limb formation, pectoral fin formation appeared normal, in the zebrafish *gpr161* mutant embryos. (Figure 2B, Figure 2 Supplement 2D).

### Hedgehog signalling is upregulated in *gpr161* mutant embryos

The morphological phenotypes observed in *gpr161* mutants are consistent with increased Hh signalling. To determine whether Hh signalling is upregulated in *gpr161* mutants, we assessed the expression of known Hh target genes in the neural tube by qRT-PCR and RNA *in situ* hybridisation. The Hh target genes *ptch2, gli1, nkx2.2b* and *nkx6.1* (Figure 3B) were all strongly upregulated, while *pax7a*, which is repressed by Hh signalling (Guner and Karlstrom, 2007), was strongly downregulated in *gpr161* mutant compared to wild type embryos (Figure 3B). RNA in situ hybridisation revealed that expression of *shha*, the major Hh ligand expressed in the neural plate was not expanded in *gpr161* mutants (Figure 3A). However, expression of *ptch2*, a direct transcriptional target of Hh signalling (Concordet et al., 1996), was expanded in *gpr161* mutants (Figure 3A), suggesting that Hh signalling is upregulated in *gpr161* mutants downstream of Shh expression. Similarly, expression of *olig2*, a marker of motor neuron induction which depends on low-level Hh activity (Dessaud et al., 2007; Park et al., 2002), as well as *nkx2.2a*, a marker for V3 interneuron progenitor cells of the lateral floorplate (Barth and Wilson, 1995; Briscoe et al., 1999), were clearly expanded in the *gpr161* mutant neural tube (Figure 3A). We note that the expansion of the low-level target *olig2* appears to be stronger than the expansion of the high-level target *nkx2.2a* (Figure 3A). Taken together, these results show that loss of Gpr161 leads to a hyperactivation of the Hh signalling pathway in zebrafish.

**Figure 3.**
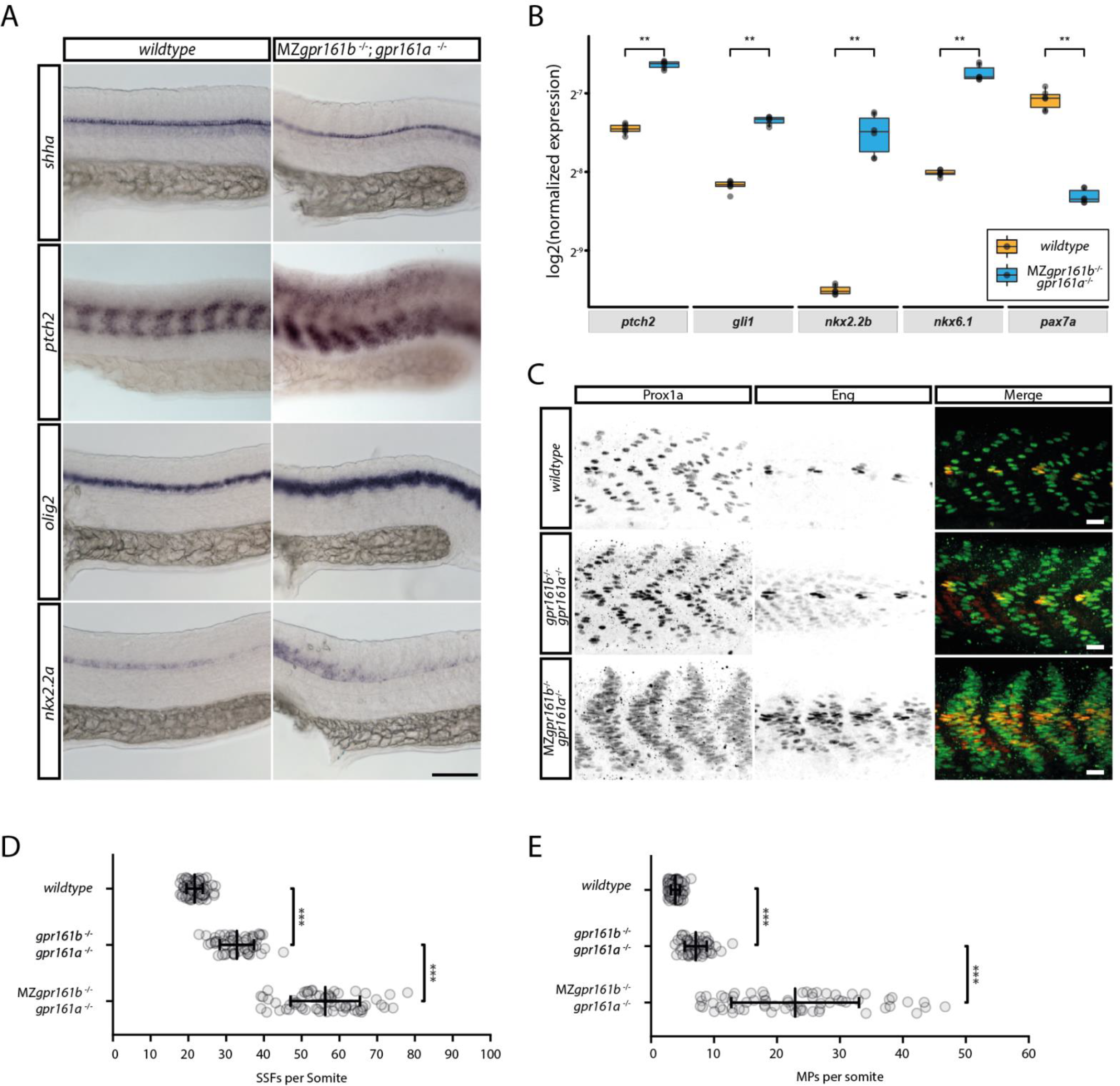
Hh signalling activity is increased in *gpr161* mutants. **(A)** RNA in situ hybridization of *shh, ptch2, olig2* and *nkx2.2a* transcripts in wildtype and MZ*gpr161b*^−/−^; *gpr161a*^−/−^ embryos fixed at 24 hpf (lateral view, scale bar: 100µm). **(B)** Transcript levels of *ptch2, gli1, nkx2.2b, nkx6.1* and *pax7a* in wildtype and MZ*gpr161b*^−/−^; *gpr161a*^−/−^ embryos at 24hpf determined by RT-qPCR (n=3)**(C)** Immunostaining of Prox1 and Eng proteins in 24 hpf zebrafish embryos reveal the number of MPs (Prox1a/Eng double positive) as well as SFFs (Prox1 positive) in wild-type, *gpr161b*^−/−^; *gpr161a*^−/−^ and MZ*gpr161b*^−/−^; *gpr161*^−/−^ embryos fixed at 24 hpf (scale bar: 20µm). Number of **(D)** SSFs and **(E)** MPs per somite in wild-type (n= 93 somites in 22 embryos), *gpr161b*^−/−^; *gpr161a*^−/−^ (n=60 somites in 20 embryos) and MZ*gpr161b*^−/−^; *gpr161a*^−/−^ (n= 66 somites in 22 embryos) embryos fixed at 24 hpf.

In the zebrafish myotome, sustained Hh signalling during gastrulation and somitogenesis stages have been shown to be required for the specification of several cell types, including Prox1 and Eng double positive muscle pioneer cells (MPs) and Prox1 positive superficial slow fibres (SSFs) (Wolff et al., 2003). While medium-to-low level Hh signalling is sufficient for the specification of SSFs, the formation of MPs requires high levels of Hh (Wolff et al., 2003). Consistent with the expansion of Hh target genes in the neural tube, *gpr161* mutants also displayed an increase in both SSFs and MPs (Figure 3C). While zygotic Gpr161 loss of function resulted in a significant increase in both SSFs (from 22 ± 2 (mean ±SD) in wt to 33 ±5 in zygotic *gpr161* mutants) and MPs (from 4 ± 1 (mean ± SD) in wt to 7 ± 2 in zygotic *gpr161* mutants), complete loss of both maternal and zygotic Gpr161 led to a stronger increase in both SSFs and MPs (56 ± 9 (mean ± s.d.) SSFs, and 23 ± 10 MPs), consistent with the requirement for sustained Hh signalling in muscle cell development (Wolff et al., 2003). These results suggest that in the somites, loss of Gpr161 results in expansion of both high and low Hh signalling targets.

### *gpr161* mutants remain sensitive to Shh

Our results suggest that although loss of Gpr161 function in zebrafish leads to increased Hh signalling activity, *gpr161* mutants display weaker phenotypes than those seen in mutants with maximal activation of Hh signalling. To determine whether Hh signalling could be further activated in *gpr161* mutants in response to Hh, we injected 50 or 100 pg *shh* mRNA, and assessed SSF and MP formation using Prox1 and Eng antibody staining as above (Figure 4A-B). In wild-type embryos, overexpression of 100 pg *shh* resulted in an approximate 1.8-fold increase in both MPs and SSFs (Figure 4B), a significant change from uninjected control embryos (ANOVA, SSFs p<0.01, MPs p<0.01). In contrast, neither 50 nor 100 pg *shh* resulted in a significant increase in the number of SSFs in *gpr161* mutant embryos (ANOVA, 50 pg p>0.9, 100 pg p>0.5), suggesting that the number of SSFs have reached a maximum level in the uninjected mutants. Injection of 100 pg *shh* did, however, give a significant 1.9-fold increase in the number of MPs in *gpr161* mutant embryos (ANOVA, p<0.001) (Figure 4B). Taken together, these results suggest that in the *gpr161* mutant, medium-to-low level Hh signalling targets are activated close to maximal levels, whereas high level targets of Hh signalling are increased, but not maximally so.

**Figure 4.**
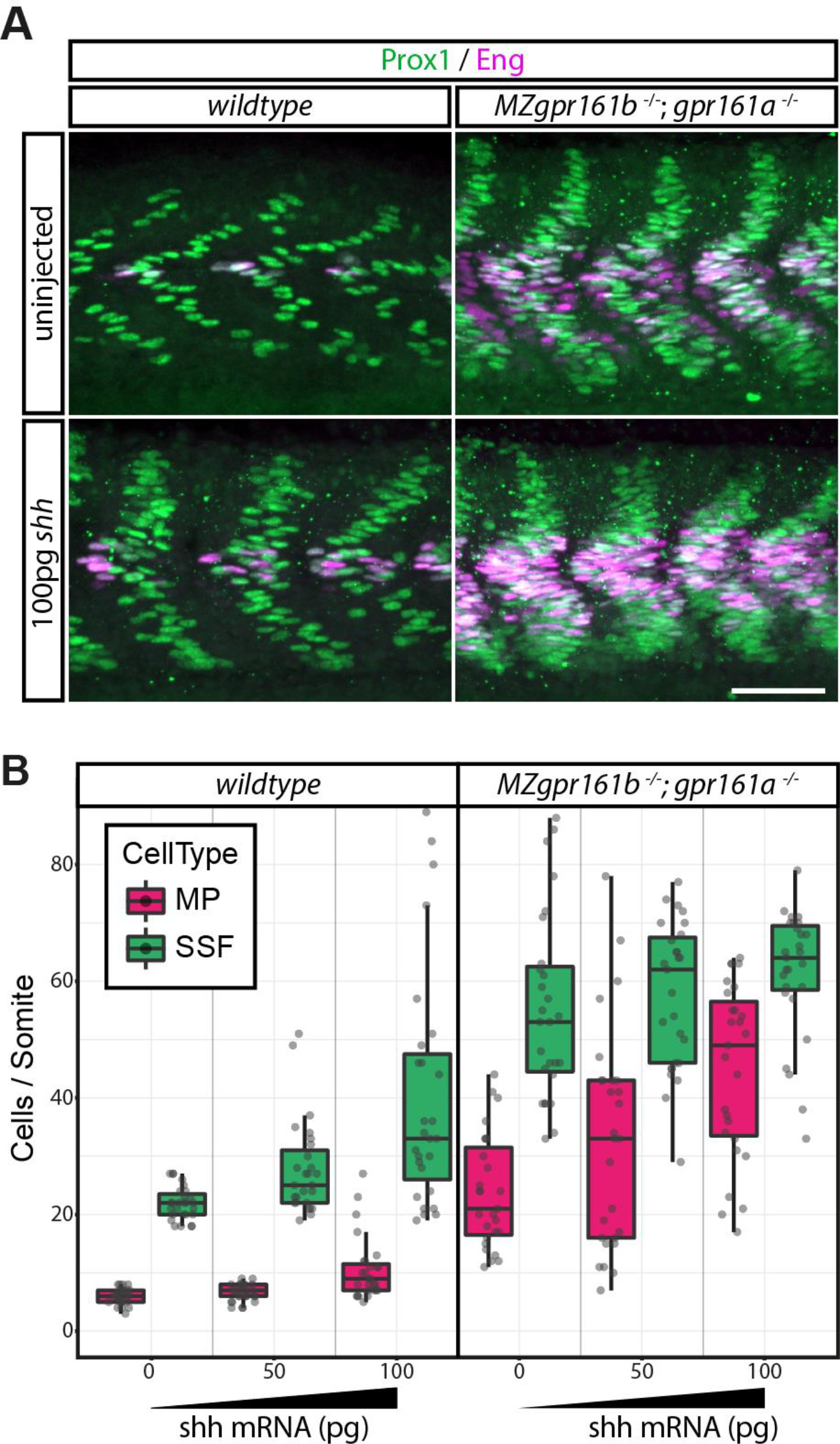
Injection of *shh* mRNA can elevate hh signalling outcomes in gpr161 mutants. **(A)** Prox1 (green)/ Eng (purple) immunostainings of wild-type and MZ*gpr161b*^−/−^; *gpr161a*^−/−^ embryos injected with 100 pg of *shha* mRNA (scale bar: 50µm). **(B)** Quantification of MPs and SSFs in wild-type and MZ*gpr161b*^−/−^; *gpr161a*^−/−^ embryos injected with increasing amounts of *shh* mRNA (for all experiments: n=27 somites in 9 embryos).

### Activation of PKA rescues *gpr161* mutant phenotypes

Murine Gpr161 was proposed to act as a constitutively active G_αs_-coupled receptor, which contributes to maintain basal levels of PKA activity to keep the Hh pathway inactive (Mukhopadhyay et al., 2013). This model predicts that the phenotypes observed in the *gpr161* mutants are due to a loss of adenylate cyclase activity, and should be rescued by artificial activation of adenylate cyclase by agents such as forskolin. Previous studies have reported that 300 µM forskolin phenocopies a complete loss of Hh signalling (Barresi et al., 2000). Consistent with this, treatment with 300 μM forskolin resulted in a near complete loss of all MPs and SSFs in wild-type as well as *gpr161* mutant embryos (Figure 5A-B), consistent with the model that hyperactivation of Hh signalling in the *gpr161* mutants is due to a reduction of PKA activity. Treatment with lower concentrations of forskolin, ranging from 500 nM to 50 µM, had only minor effects on somite development of wild-type embryos, but resulted in a significant degree of rescue of the *gpr161* mutant phenotype (Figure 5A-B). We note that MPs, which require high levels of Hh signalling, were more sensitive to forskolin concentrations than SSFs, which require low levels of Hh.

**Figure 5.**
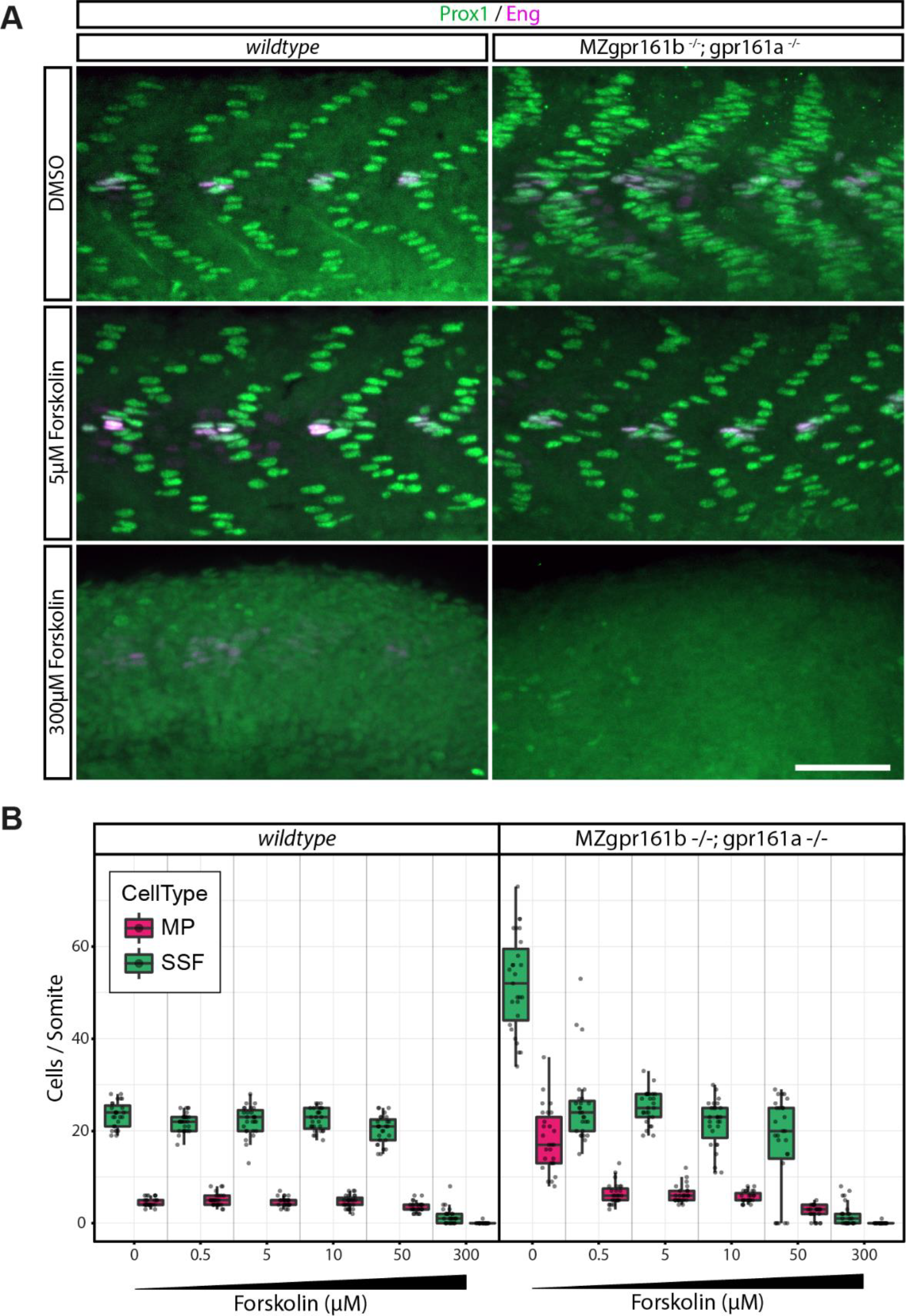
Forskolin treatments can rescue the gpr161 phenotype. **(A)** Prox1 (green)/ Eng (purple) immunostainings of wild-type and MZ*gpr161b*^−/−^; *gpr161a*^−/−^ embryos treated with increasing concentrations of forskolin (scale bar: 50µm). **(B)** Quantification of MPs and SSFs in wild-type and MZ*gpr161b*^−/−^; *gpr161a*^−/−^ embryos treated with increasing concentrations of forskolin (for each experiment, n=27 somites in 9 embryos).

### Distinct contributions of Gpr161 to basal repression of Hh signalling in different tissues

To determine the epistatic relationship between Smo and Gpr161 in zebrafish, we used the zebrafish *smo* null allele *hi1640* (Chen et al., 2001) to generate zebrafish *gpr161b*^−/−^; *gpr161a*^−/−^; *smo*^−/−^ triple mutants, and assessed Hh signalling activity using Prox1/Eng staining and neural tube markers, as above. Zebrafish *gpr161b*^−/−^; *gpr161a*^−/−^; *smo*^−/−^ triple homozygous mutant embryos showed a morphological phenotype, as well as an expansion of both SSFs and MPs, that were indistinguishable from *gpr161b*^−/−^; *gpr161a*^−/−^ double homozygous mutant embryos (Figure 6A-B, Figure 6 Supplement 1), consistent with the reported phenotypes in the mouse *Gpr161;Smo* double mutants (Mukhopadhyay et al., 2013). Further, the expansion of low-level Hh target-genes *ptch2* and *olig2* was similarly found to be independent of Smo function (Figure 6C). In contrast, the double mutant *gpr161*^−/−^; *gpr161a*^−/−^; *smo*^−/−^ triple mutant embryos showed no detectable *nkx2.2a* expression, suggesting that in the neural tube, though not in the somites, high level Hh target gene expression is dependent on Smo function in the absence of Gpr161.

**Figure 6.**
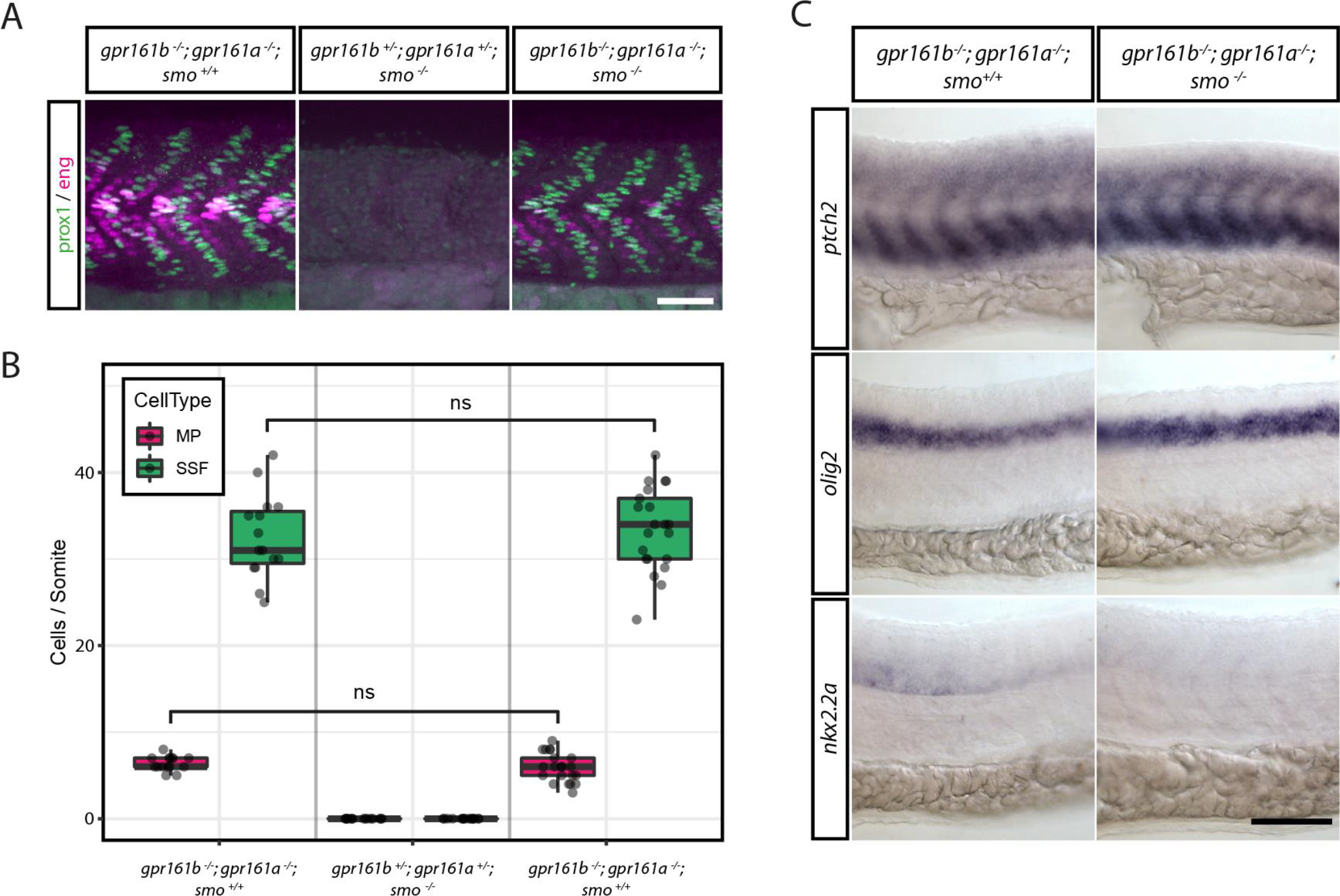
The requirement for Smo during high-level Hh signalling in *gpr161* mutants is tissue-specific. **(A)** Prox1 (green)/ Eng (purple) immunostainings of wildtype and MZgpr161b^−/−^; gpr161a^−/−^ embryos at 24 hpf after DMSO or 200µM Cyclopamine treatment (scale bar: 50µm). **(B)** Quantification of MPs and SSFs from experiment presented in A (*gpr161b*^−/−^; gpr161a^−/−^; *smo*^+/+^ and *gpr161b*^+/−^; *gpr161a*^+/−^; *smo*^−/−^: n=15 somites in 5 embryos; *gpr161b*^−/−^; *gpr161a*^−/−^; *smo*^−/−^: n=21 somites in 7 embryos; all others: n=27 somites in 9 embryos). **(C)** RNA in situ hybridization of *ptch2, olig2* and *nkx2.2a* transcripts in *gpr161b*^−/−^; gpr161a^−/−^; smo^+/+^ and *gpr161b*^−/−^; *gpr161a*^−/−^; *smo*^−/−^ embryos fixed at 24 hpf (lateral view, scale bar: 100µm).

## Discussion

Hh signalling is tightly controlled at multiple levels in order to accurately translate morphogen gradients into graded transcriptional responses mediated by the Gli transcription factors. PKA both promotes formation of the Gli repressor forms and inhibits Gli activator forms, thereby controlling sensitivity to Hh ligands, and providing a filter for Hh signalling strength. How PKA activity is fine-tuned in Hh signalling is not completely understood. GPCRs such as Gpr161 have been shown to impact on Hh signalling by regulating PKA via the regulation of adenylate cyclase activity. To extend our understanding of how GPCRs regulate Hh signalling, we generated zebrafish *gpr161* mutants and analysed Hh-dependent signalling in the neural tube and the myotome.

The zebrafish genome contains two conserved Gpr161 orthologues, Gpr161a and Gpr161b. We have generated mutants for both orthologues, and show that Gpr161a and Gpr161b act redundantly in early zebrafish development. Zebrafish mutants lacking Gpr161 function show an expansion of Hh target gene expression both in the neural tube and in the somites, and develop severe craniofacial defects that are similar to those described for the *ptc1*^−/−^; *ptc2*^−/−^ double mutants (Koudijs et al., 2008). These results confirm that the role of Gpr161 as a modulator of Hh signalling is conserved in the vertebrate lineage.

Gpr161 was proposed to contribute to the basal Hh repression machinery by activating G_αs_ and adenylate cyclase, resulting in activation of PKA, and overexpression of murine Gpr161 was shown to lead to a general increase in intracellular cAMP levels (Mukhopadhyay et al., 2013). However, direct evidence for reduced cAMP production in Gpr161 mutant cells is hampered by the difficulties in measuring physiological cAMP levels in specific subcellular compartments such as the primary cilium. We have taken advantage of the zebrafish embryo’s amenability to pharmacological treatments to further probe the mechanism of Gpr161 action. Consistent with the model that loss of Gpr161 leads to lowered cAMP levels and reduced PKA activity, we found that treatment with the cAMP elevating agent forskolin fully rescued muscle cell specification in the *gpr161* mutant embryos. Interestingly, a 100-fold concentration range (0.5-50 µM) of forskolin gave very similar near-complete rescue of both mutant morphology and molecular read-outs of both high and low level Hh target gene expression in the somites, suggesting that additional mechanisms may be in place to ensure appropriate regulation of PKA activity in the presence of excess cAMP. Previous studies have reported that loss of Gpr161 may also affect other signalling pathways in addition to Hh signalling (Li et al., 2015; Mukhopadhyay et al., 2013). While we cannot rule out a role for Gpr161 in processes not related to Hh signalling, our results suggest that loss of Gpr161 function can be fully compensated by artificial activation of adenylate cyclase.

A comparison of phenotypes suggests that the upregulation of Hh signalling in zebrafish *gpr161* mutants is less severe than that seen in *ptc1*^−/−^; *ptc2*^−/−^ mutant embryos (Koudijs et al., 2005). Similarly, injection of a dominant-negative form of PKA resulted in an apparently stronger increase in Hh-dependent muscle cell specification in the myotome than could observed in the *gpr161* mutants (Zhao et al., 2016). This is consistent with data obtained in mice, where a loss of PKA or G_αs_ leads to more severe ventralisation of the neural tube than what is observed in the *Gpr161* mutants (Mukhopadhyay et al., 2013; Pusapati et al., 2018; Regard et al., 2013; Tuson et al., 2011). Further supporting the idea that Hh signalling is not maximally activated in the *gpr161* mutants, we find that injection of *shha* mRNA can further increase high, but not low, level Hh targets in the somites of *gpr161* mutants. Thus we conclude that whereas low level Hh signalling is maximally active in the *gpr161* mutants, additional mechanisms contribute to PKA activation to control high level Hh signalling in the absence of Gpr161 function.

Importantly, we also show that low level Hh signalling in the neural tube of *gpr161* mutants is independent of Smo, whereas expression of the high level Hh target gene *nkx2.2a* clearly requires Smo function. Our results are consistent with the model that G_αs_-coupling and activation by Gpr161 is one of several mechanisms that contribute to the mobilisation of compartmentalised cAMP to repress Hh target activation, and that the reduction in cAMP levels caused by loss of Gpr161 is sufficient to cause constitutive, Smo-independent activation of low, but not high level Hh signalling. This result provides genetic evidence to support the model based on results from pharmacological inhibition of Smo in mammalian cell culture (Pusapati et al., 2018). Similar experiments were performed in the mouse *Gpr161* mutant, with the conclusion that Gpr161 is largely epistatic to Smo (Mukhopadhyay et al., 2013). These authors do however note that the expression of high level Hh target genes, such as Nkx2.2 and FoxA2, is reduced in *Smo; Gpr161* double mutants compared to *Gpr161* single mutants. One possibility is that this difference is due to the different assays used to assess Hh target gene expression in mouse and zebrafish neural tubes. Whereas Mukhopadhyay and colleagues used immunohistochemistry to detect Hh target gene expression (Mukhopadhyay et al., 2013), our results are based on chromogenic in situ hybridisation, a far less sensitive assay. Thus, we can not rule out that some low level *nkx2.2a* expression persists in the zebrafish triple mutants. Another possibility is that there could be species-specific differences in the roles of GliR and GliA and/or cAMP levels, or alternatively, Gpr161 may make a relatively larger contribution to cAMP levels in the zebrafish neural tube compared to mouse. Interestingly, we do observe tissue-specific differences in zebrafish in our epistasis experiments. In the *gpr161* mutant myotome, both high and low level Hh signalling outcomes are independent of Smo, suggesting that in the somites Gpr161 is completely epistatic to Smo. We suggest that in the neural tube, additional unknown factors make significant contributions to promote basal cAMP levels, whereas in the zebrafish myotome, Gpr161 alone may account for the largest part of the basal repression machinery. Thus, distinct combinations of GPCRs in different cell types can contribute to complex and tissue-specific regulation of Hh signalling. The identification of these additional GPCRs, as well as other factors that control PKA activity downstream of adenylate cyclases, will be required to understand how Hh signalling is fine-tuned to orchestrate the great variety of Hh-dependent biological processes in a cell type specific manner.

## Materials and Methods

### CRISPR/cas9 genome editing and genotyping

Guide RNAs for CRISPR/cas9 mediated knockout of both *gpr161a* (ENSDART00000151311.2) and *gpr161b* (ENSDART00000078051.6) were designed using the ChopChop web tool (Montague et al., 2014) and synthesised as described previously (Huang et al., 2014). Embryos were injected with 50 pg of gene specific sgRNA and 300pg of *cas9* mRNA at 1-cell stage. F0 founder fish were identified by T7 Endonuclease I digests of gene-specific PCR products from pooled genomic DNA obtained from F1 offspring, following the manufacturer’s protocol (NEB, #M0302L). While the *gpr161a*^*ml200*^ allele harbors an 8bp deletion, the introduced mutation in *gpr161*^*ml201*^ leads to a 6bp insertion, which were identified by running out gene specific PCR products (see Table 2) on 4% agarose gels. Genotyping of the *smo*^*hi1640*^ allele was performed as described previously (Chung and Stainier, 2008).

**Table 1.**
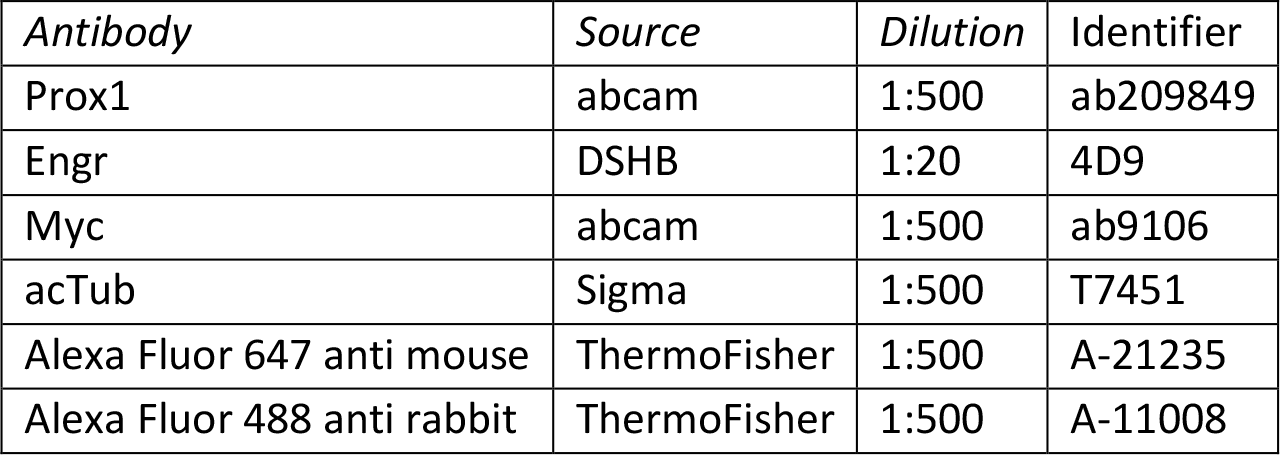
List of Antibodies

**Table 2.**
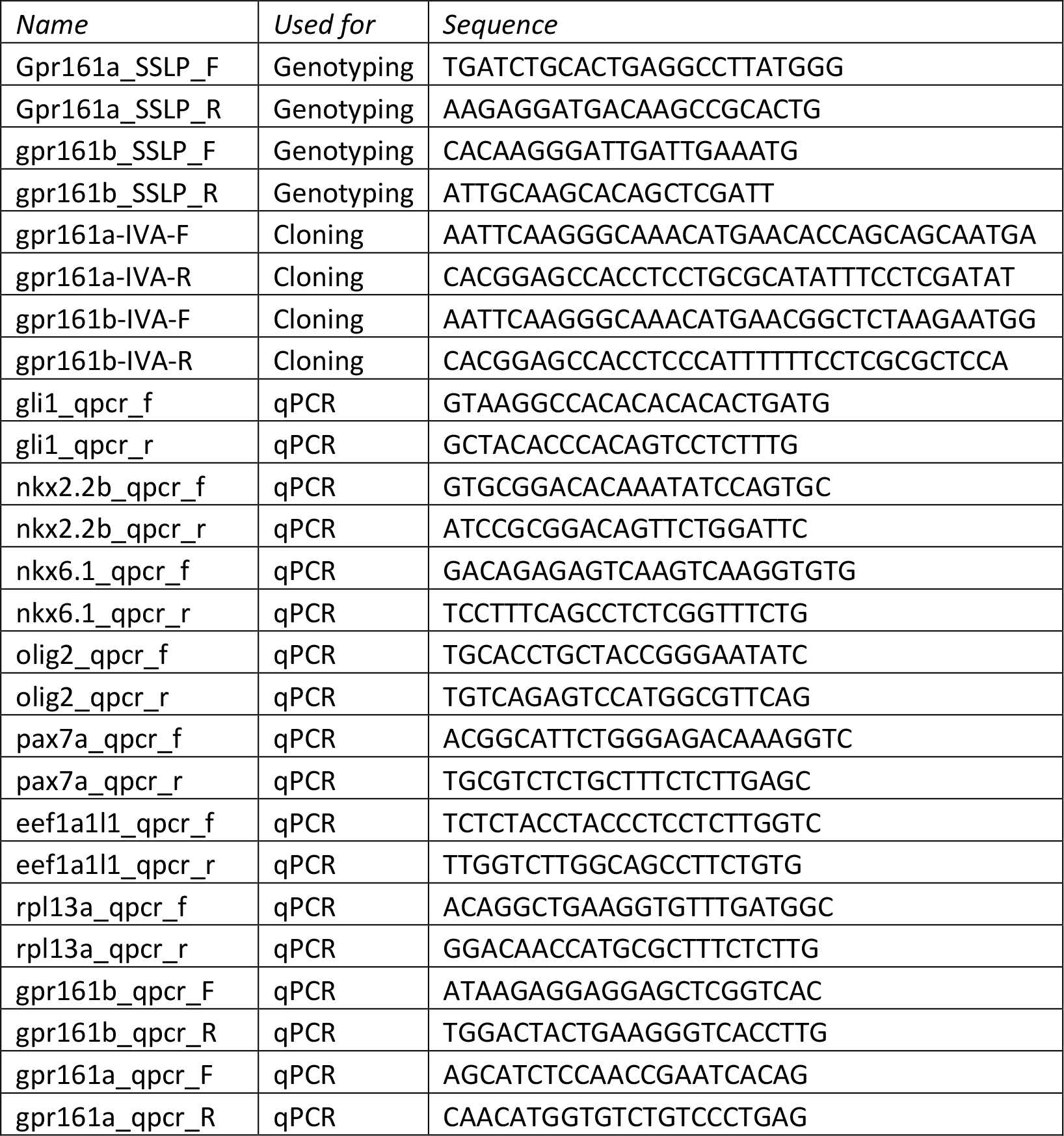
List of Primers

### Zebrafish lines and husbandry

All zebrafish lines including SAT wildtype strains were kept at 28°C according to standard protocols.

MZ*gpr161b*^−/−^; *gpr161a*^−/−^ embryos were obtained by performing in-crosses of a *gpr161b*^−/−^; *gpr161a*^+/−^ line. Embryos were raised at 28°C and staged by morphology (Kimmel et al., 1995).

All experimental protocols concerning zebrafish were approved by the Austrian Ministry for Science and Research (BMWFW-66.008/0016-WF/V/3b/2016, BMBWF-66.008/0015-V/3b/2018), and experiments were carried out in accordance with approved guidelines.

### Construction of plasmids

To generate expression-vectors for *gpr161a* and *gpr161b* the coding sequence of both genes were amplified with overlapping primers (see Table 2) using homemade PfuX7 polymerase (Nørholm, 2010) and fused in-frame to a Myc-Tag into pCS2+ by *in vivo* Assembly (IVA) cloning (García-Nafría et al., 2016).

### Quantitative (q) RT-PCR

RNA was isolated from zebrafish embryos with Trizol (Ambion) following the manufacturer’s instructions. RNA integrity was checked by agarose gel electrophoresis and the concentration was measured using a Nanodrop 2000c (Thermo) spectrophotometer.

Complementary DNA was transcribed from equal amounts of dsDNase-treated total RNA using the Maxima RT kit for qPCR (Thermo) with dsDNase according to the manufacturer’s instructions.

RT-qPCRs were performed using 5x HOT FIREPol EvaGreen qPCR Supermix (Solis Biodyne) and contained each primer at 250nM and cDNA corresponding to a total RNA amount of 15 ng for pooled embryos or 5 ng for single embryos. PCRs were run on a CFX96 Connect (BioRad) under following conditions: 12 min 95°C, 40 cycles of 95°C for 30s, 60°C for 30s and 72°C for 20s. Melt curves were recorded from 65°C to 95°C in 0.5°C increments. Data was acquired using CFX Manager 3.1 (BioRad) and exported as RDML files for processing.

Data analysis was performed in R version 3.4.4. Fluorescence data were imported using the package RDML (Rödiger et al., 2017) and amplification curves fitted using the ‘cm3’ model (Carr and Moore, 2012) implemented in the package qpcR (Ritz and Spiess, 2008). The first derivative (d0) of the model was used as expression value. Expression values for genes of interest were normalised using the geometric mean of the expression values of the reference genes *eef1aa* and *rpl13*.

### Whole-mount in situ hybridisation

In situ hybridisation was performed following standard protocols. DIG-labelled antisense probes were made for *shha* (Krauss et al., 1993), *ptch2* (Concordet et al., 1996), *olig2* (Park et al., 2002) and *nkx2.2a* (Barth and Wilson, 1995).

### Immunohistochemistry

Embryos were fixed in 4% paraformaldehyde at room temperature for 3h, then washed in PBS-Triton (PBS + 0.3% Triton X-100). After 1h of incubation in blocking solution (PBS-Triton, 4% BSA, 0.02% NaN_3_) at 4°C, primary antibodies diluted in blocking solution were added and left over night for incubation at 4°C. After subsequent washes in PBS-Triton embryos were incubated with appropriate Alexa conjugated secondary antibodies over night at 4°C. Again, Embryos were washed several times in PBS-Triton and mounted in Mowiol embedding medium for imaging. For a list of the antibodies used in this study, see Supplementary Table 1. All stainings were imaged using a Zeiss Axio Observer.Z1 microscope equipped with a Yokogawa CSU-X1 spinning disc confocal unit using 25x, or 63x water-immersion lenses.

### Chemical treatments

For chemical treatments embryos were dechorionated at 50% epiboly and transferred to agar-coated 35mm dishes containing forskolin (Biomol) at final concentrations between 0.5 and 300 µM in 1% DMSO. Control experiments were performed simultaneously in 1% DMSO. All embryos were treated until 24 hpf.

### Microinjection

mRNA was synthesised using the HiScribe SP6 RNA Synthesis Kit (NEB) and capped using the Vaccinia Capping System (NEB) following the protocols provided by the manufacturer. Embryos were injected at one-cell stage. Injected volumes are indicated in the respective figures.

### Light Histology

Three day old (72 hpf) wild type and *gpr161* mutant embryos were fixed in 2.5% glutaraldehyde in 0.01M sodium cacodylate buffer for two hours, washed in buffer, dehydrated in an increasing acetone series and embedded in EMBed 812 epoxy resin. After polymerisation for 48 hours at 60°C, embryos were cut serially with an Autocut 5020 (Reichert, Austria) and a Diatome Butler knife (Diatome, Switzerland). 2 µm thick serial sections were stained according to Richardson (Richardson et al., 1960) for 10 minutes, washed, and mounted in cedarwood oil. Images were taken with a Leica DM5000B microscope using a Leica DFC 490 digital camera and Leica application suite v. 4.8 (Leica, Germany).

### Electron microscopy

Embryos were fixed with 2.5% glutaraldehyde in 0.01M sodium cacodylate buffer containing 5% sucrose at 4°C for two hours. After washing in cacodylate buffer, specimens were post fixed in reduced osmium (2% osmium tetroxide and 3% potassium ferrocyanide in 0.1M cacodylate buffer) for two hours at 4°C, dehydrated in an ethanol series, critical point dried with an EMS 850 CPD (Electron Microscopy Services, Germany), mounted and 20 nm gold sputtered with a CCU-010 sputter coater (Safematic, Switzerland), and examined with a DSM950 scanning electron microscope (Zeiss, Germany). Images were taken with a Pentax digital camera and PK_Tether 0.7.0 free software.

### Data Presentation and Analysis

All data presented in this study were analyzed with R using the RStudio integrated development environment and plotted using the “ggplot2” package (Rstudio Team, 2016; Wickham, 2016).

Statistical significance of differences in expression levels between groups were calculated on at least three biological replicates with the Wilcoxon rank sum test, corrected for multiple comparisons using the Benjamini-Hochberg FDR method.

Statistical significance of a difference in MP or SSF numbers between groups was determined using one-way ANOVA corrected for unequal variances and the Games-Howell post-hoc test for pairwise comparison as implemented in the “userfriendlyscience” package (Peters, 2017). P values are indicated as follows: * p<0.05; ** p<0.01; p<0.001; ns not significant. Sample sizes (N) are given in the respective figure legends.

## Acknowledgements

We are grateful to Dzenana Tufegdzic for fish care, and to Dirk Meyer, Robin Kimmel and Kathi Klee for helpful discussions and/or comments on the manuscript. This work was supported by funding from the Austrian Science Fund (FWF) and the Tyrolean Science Fund (TWF) (FWF P27338 (to P.A.), FWF P30441 (to E.S.), TWF 236277 (to D.R.)).

## Competing interests

The authors declare they have no competing interests.

---

**Figure 1-Supplement 1.**
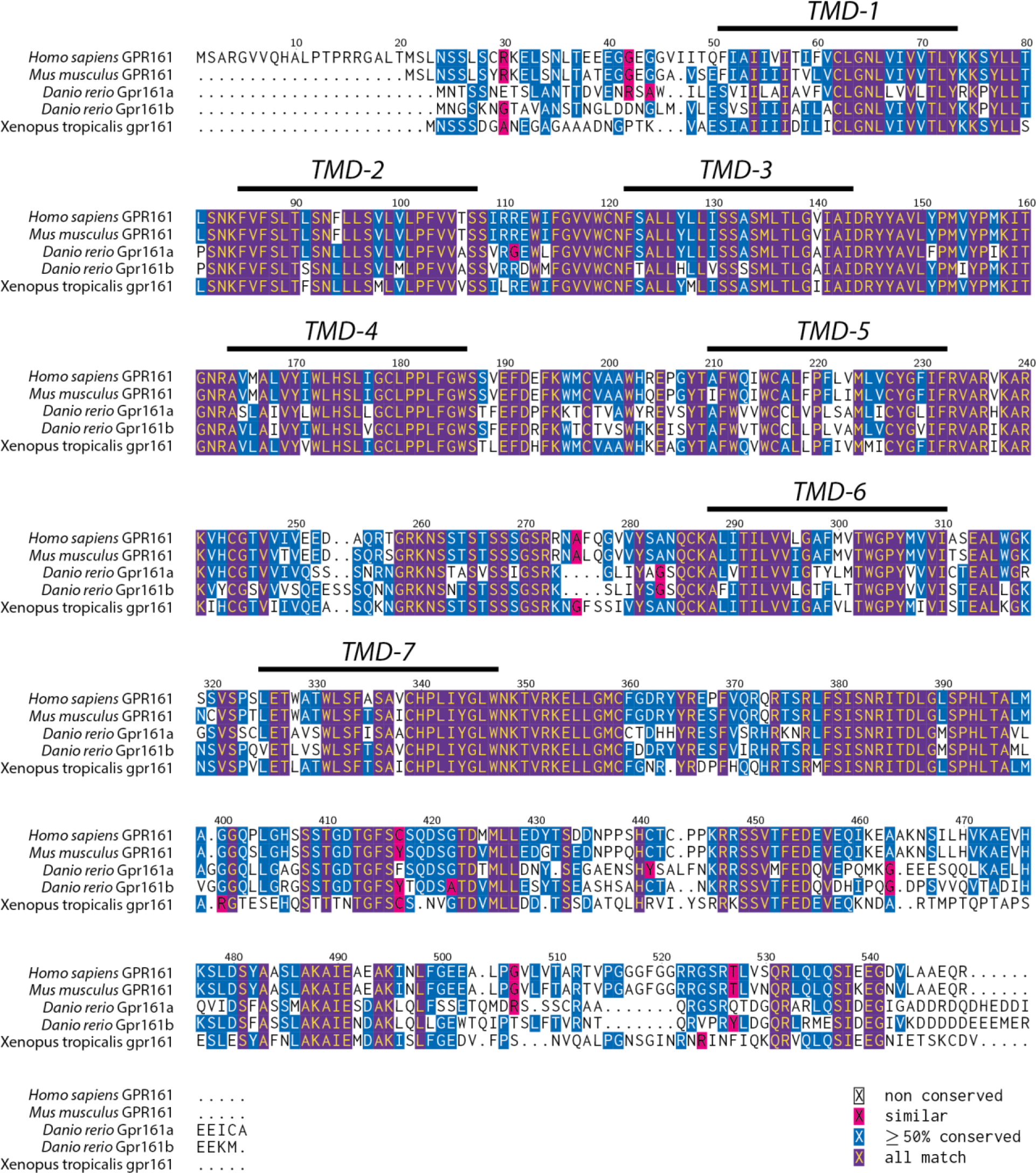
Multiple Sequence Alignment. The sequences of the *Homo sapiens, Mus musculus, Danio rerio* and *Xenopus tropicalis* Gpr161 proteins were aligned using MUSCLE (Edgar, 2004). Transmembrane domains (TMDs) and conserved residues are indicated.

**Figure 1-Supplement 2.**
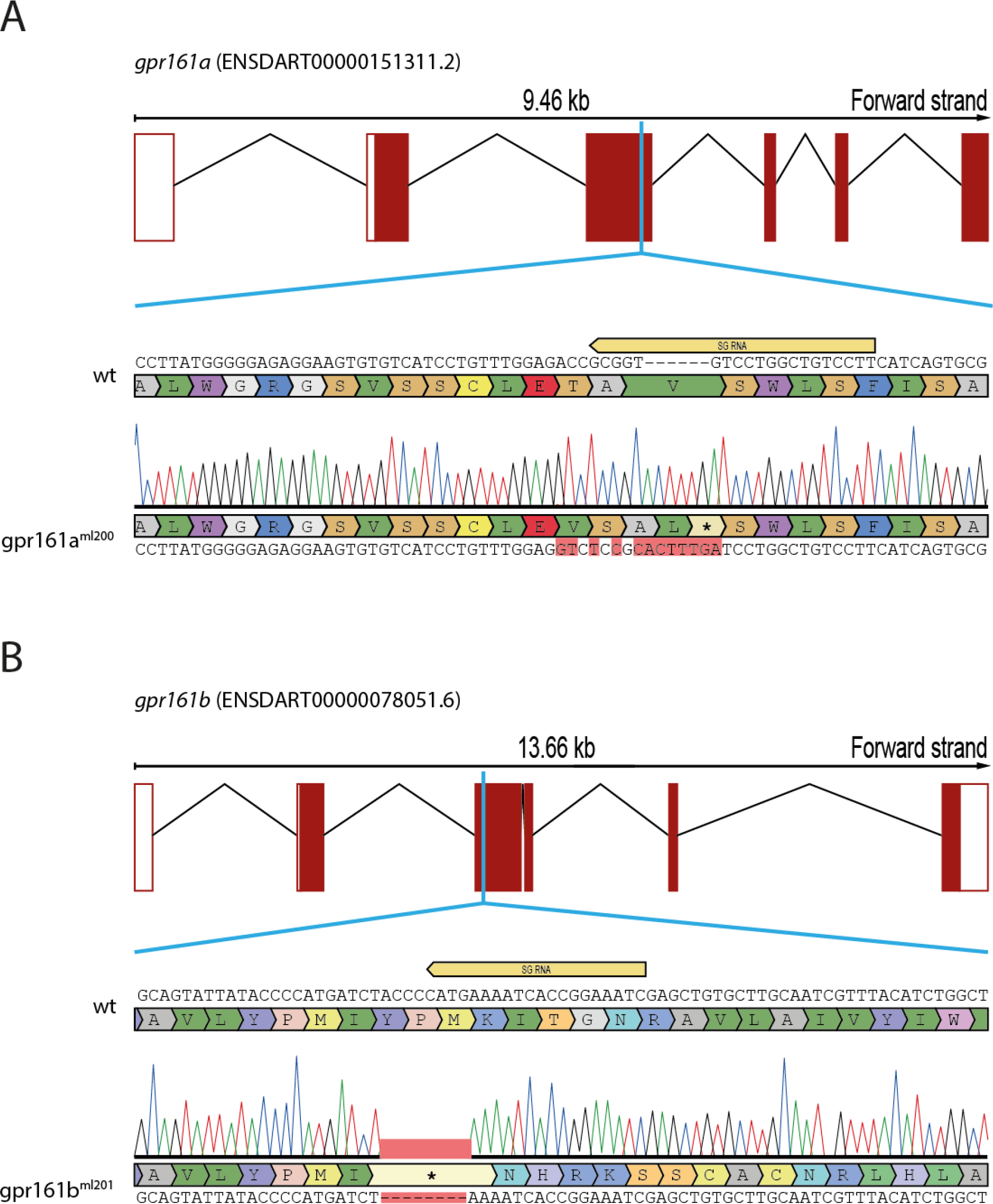
CRISPR knock out strategy. Schematic representation of the gene structure of *gpr161a* **(A)** and *gpr161b* **(B)** indicating the position of CRISPR sgRNA recognition site and a sequence alignment of the obtained mutant alleles.

**Figure 2-Supplement 1.**
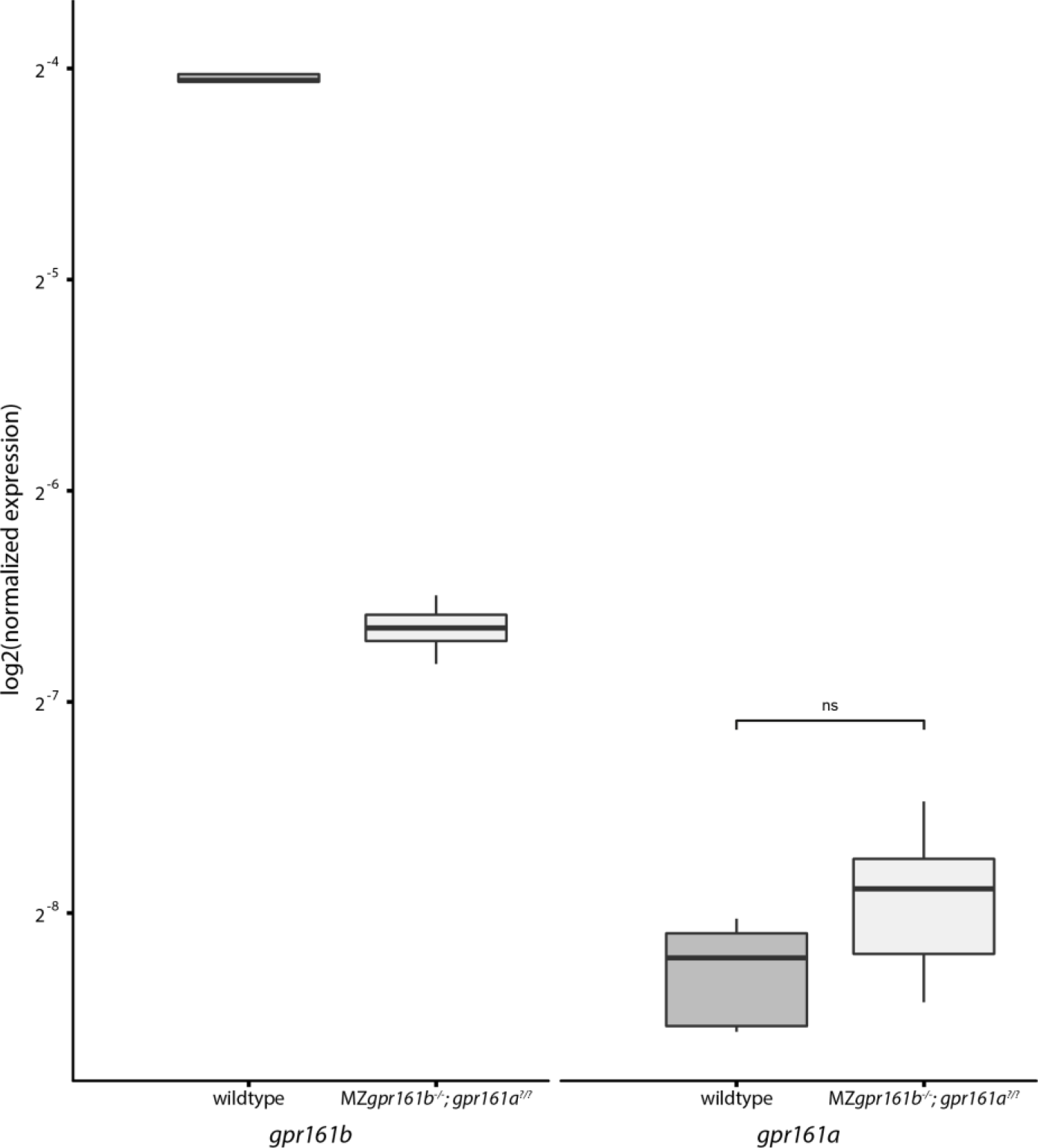
Maternal loss of *gpr161b* does not lead to genetic compensation by *gpr161a.* Transcript levels of *gpr161a* and *gpr161b* in single embryos at 2-cell stage as determined by qPCR (n=6)

**Figure 2-Supplement 2.**
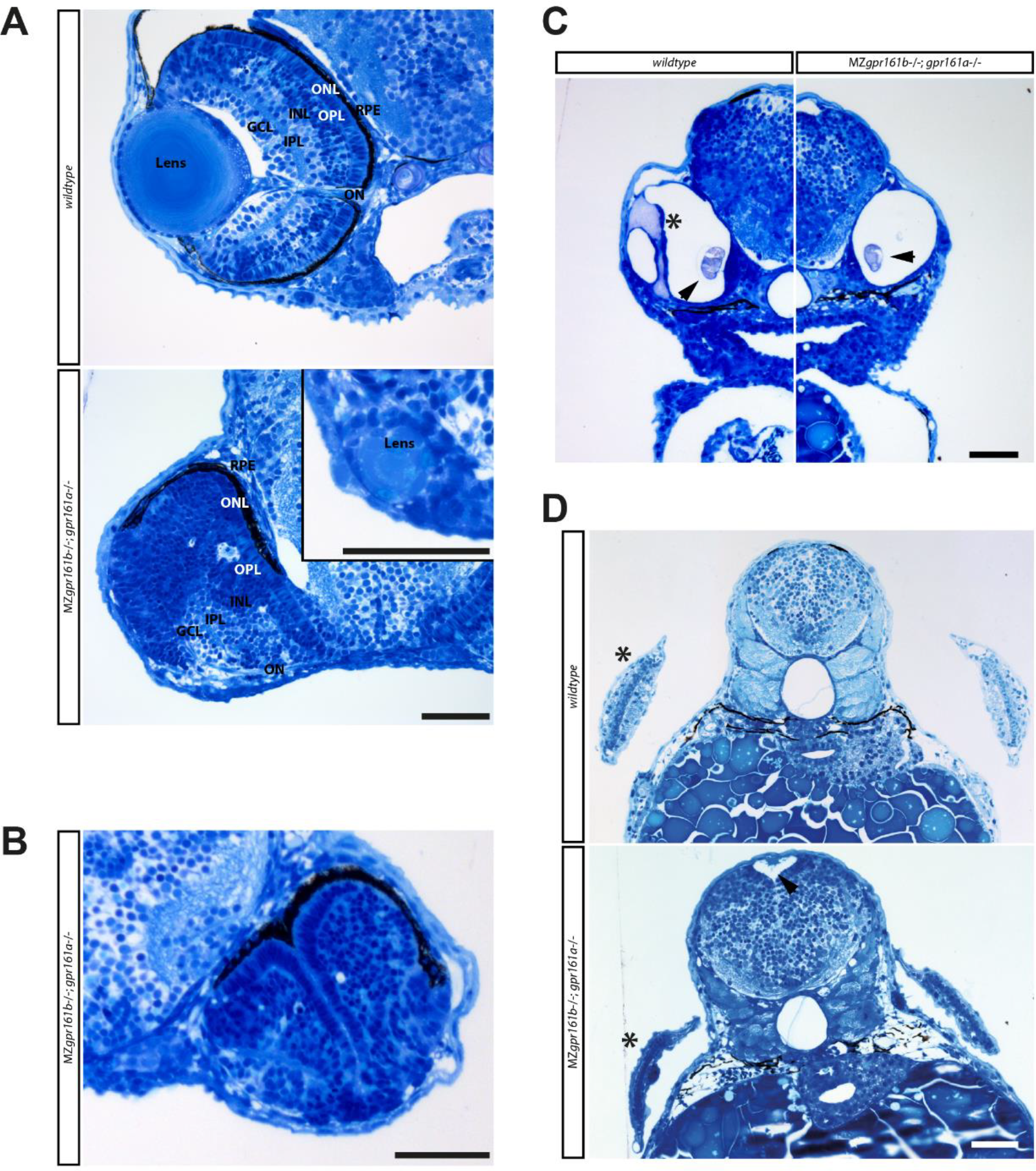
Severe morphological defects in *gpr161* mutant embryos at 72 hpf. **(A)** Transverse section through the embryonic eye at 72 hpf, Retinal layers are indicated as follows: RPE – retinal pigment epithelium, ONL – outer nuclear layer, OPL - outer plexiform layer, INL – inner nuclear layer, IPL inner plexiform layer, GCL ganglion cell layer. Inset shows remnant of a forming lens from a different section of the same *gpr161* mutant embryo. **(B)** Transverse section through the embryonic eye of a 72 hpf *gpr161* mutant embryo shows a conspicuous invagination of the outer retinal layers. **(C)** Transverse section through the otic capsule of wildtype and *gpr161* mutant embryos at 72 hpf. Asterisk highlights central canal and dorsolateral septum which are missing in *gpr161* mutant embryos. Arrowhead shows otholits. **(D)** Transverse section through the hindbrain region of wildtype and *gpr161* mutant embryos at 72 hpf. Asterisks highlight presence of pectoral fins. Arrowhead shows enlarged brain ventricle. An increase in hindbrain diameter can be noted. (all scale bars: 50µm)

**Figure 6-Supplement 1.**
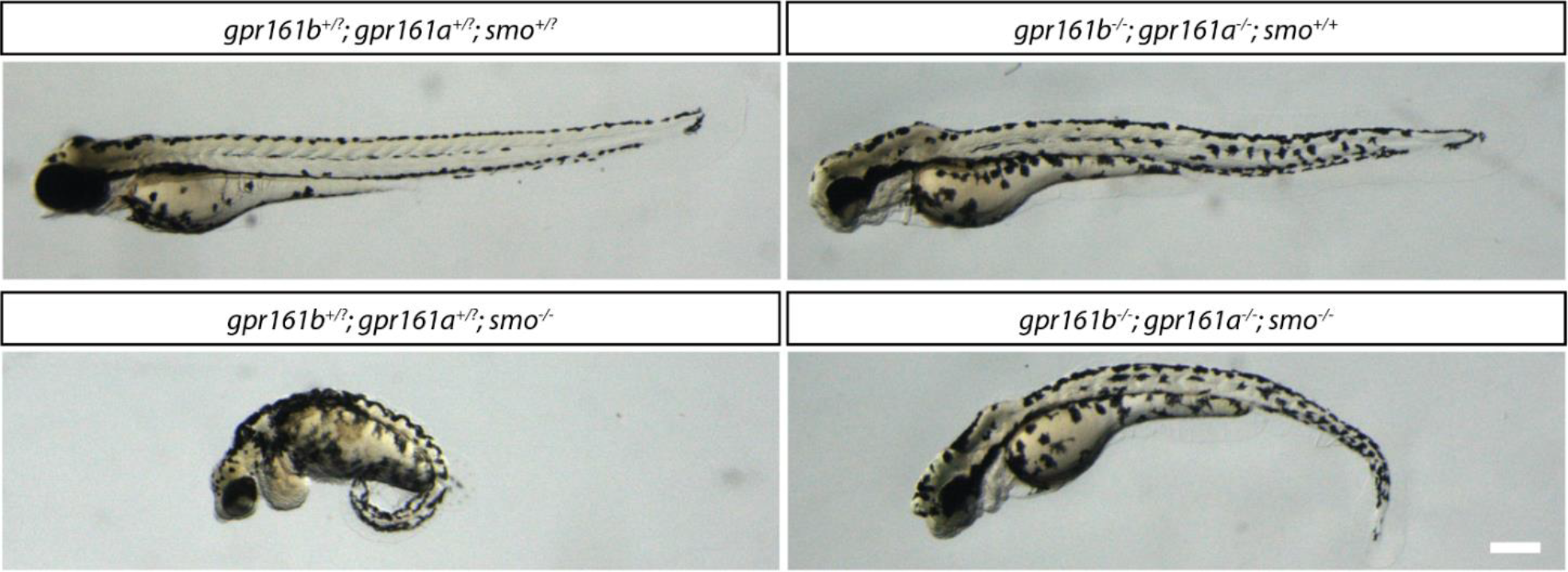
Morphological phenotypes of *gpr161b*^−/−^; *gpr161a*^−/−^ mutants are independent of Smo. Lateral view of embryos from a *gpr161b*^+/−^; *gpr161a*^+/−^; *smo*^+/−^ incross at 72hpf (scale bar: 100µm).

